# Divergent mitochondrial and nuclear OXPHOS genes are candidates for genetic incompatibilities in *Ficedula* Flycatchers

**DOI:** 10.1101/588756

**Authors:** Eva van der heijden, S. Eryn McFarlane, Tom van der Valk, Anna Qvarnström

## Abstract

Hybrid dysfunction is an important source of reproductive isolation between emerging species. Bateson-Dobzhansky-Muller incompatibilities are theoretically well-recognized as the underlying cause of low hybrid dysfunction. However, especially in wild populations, little empirical evidence exists for which genes are involved in such incompatibilities. The relative role of ecological divergence in causing the build-up of genetic incompatibilities in relation to other processes such as genomic conflict therefore remains largely unknown. Genes involved in energy metabolism are potential candidates for genetic incompatibilities, since energy metabolism depends on co-expression of mitochondrial DNA (mtDNA) and nuclear DNA (nDNA) leading to mitonuclear coadaptation. When mitochondrial and nuclear genes lacking a co-evolutionary history appear together in hybrids, incompatibilities could arise. *Ficedula* flycatcher F1 hybrids have a higher resting metabolic rate (RMR) compared to the parental species, which could be a sign of genetic incompatibilities between energy metabolism genes that diverged in response to environmental differences while the species were in allopatry. Based on sequences of 15 mitochondrial genes of 264 individuals, we show that the two species have divergent mtDNA caused by the build-up of mainly synonymous mutations and a few non-synonymous mutations. Pied flycatcher mitogenomes show evidence of non-neutrality, indicating a selective sweep or population expansion. There is little variation in the nuclear OXPHOS-related proteins and no significant deviation from neutrality, however, specific codon identified sites might be under positive selection in both mitochondrial and nuclear genes encoding OXPHOS proteins for complex I and III. Taken together, these diverged mitonuclear genes therefore constitute possible candidates underlying, at least part of the genetic incompatibilities that cause hybrid dysfunction in crosses between collared and pied flycatchers.

## Introduction

Sterility or inviability is often observed in the F1 hybrids of genetically diverged populations. Bateson (1909), Dobzhansky (1936) and Muller (1940; 1942) outlined theoretical models explaining the evolution of such genetic incompatibilities in hybrids. They proposed a two-loci system with interacting genes where alternative new mutations can go to fixation in geographically separated populations without causing incompatibilities within any one of the populations (i.e. Bateson-Dobzhansky-Muller incompatibilities; BDMI). However, when the two populations come into secondary contact and interbreed, these alternative alleles can cause hybrid dysfunction as a result of epistasis. Empirical evidence for BDMI has been found in model species such as *Mimulus* (Fishman & Willis 2001) and *Drosophila* (Brideau et al 2006),but general processes that lead to BDMIs are not well described in wild systems. To what extent ecological divergence, rather than other processes such as genomic conflict, is causing the build-up of genetic incompatibilities therefore remains largely unknown.

One set of genes that are potentially interesting candidates for BDMI are those directly involved in energy metabolism(Burton & Barreto 2012, Gershoni et al 2009). The fitness of an organism depends upon efficient energy metabolism under the environmental conditions experienced (e.g. (Bozinovic et al 2011, Mishmar et al 2003). Thus local adaptations to climate can cause rapid divergence between populations (Qvarnström et al 2016). Energy production is regulated by the mitochondria via the oxidative phosphorylation pathway (OXPHOS). Mitochondrial DNA (mtDNA) codes in most organisms for 13 proteins and several non-coding RNAs (transfer RNA, ribosomal RNA). OXPHOS is built up of protein complexes composed of both mitochondrial and nuclear encoded proteins and thus mtDNA and nuclear (nDNA) products strongly interact in this pathway(Smeitink et al 2004). This results in selection for coadaptation between these two genetic systems (Gershoni et al 2009, Willett & Burton 2001). MtDNA mutates quickly (Brown et al 1979) and usually does not recombine, which, in combination with a small effective population size, may lead to fixation of slightly deleterious mutations. Coevolution can then lead to compensatory changes in the interacting nDNA (Rand et al 2004). If energy metabolism in two populations has adapted to different climates (e.g. changes in mtDNA and the interacting nDNA), hybridisation can result in incompatibilities between the OXPHOS proteins in the F1 offspring i.e. BDMIs.

Natural hybrid zones provide good opportunities to study the possible effect of divergent energy metabolism on postzygotic isolation. *Ficedula hypoleuca* and *Ficedula albicollis* (pied and collared flycatchers, respectively) are two species of small migratory birds that diverged less than one million years ago (Nadachowska-Brzyska et al 2013) and are frequently used as models for research on ecology and evolution (Qvarnström et al 2010). Pied flycatchers are present mostly in the north of Europe, whereas collared flycatchers have a more southern breeding distribution (Figure 1). In the 1960’s, collared flycatchers colonized the Swedish island Öland that was already inhabited by pied flycatchers (Qvarnström et al 2010), thus leading to secondary contact between the two species. Divergent climate adaptation seems to play an important role in mitigating the effects of competitive interactions between the two species of flycatchers (Qvarnström et al 2016). Collared flycatchers displace pied flycatchers from the preferred breeding sites (Vallin et al 2012), but appear to be constrained to a narrower niche use, whereas pied flycatchers are less affected by a mismatch between food abundance and nestling growth (Qvarnström et al 2009, Sirkiä et al 2018). This robustness might be explained by higher plasticity in metabolic rate of pied flycatchers, as pied flycatcher nestlings have a lower resting metabolic rate (RMR) at higher temperatures (associated with low food availability) and increased RMR in response to relaxed sibling competition when compared to collared flycatchers (McFarlane et al 2018). The flexible RMR response to environmental conditions observed in pied flycatchers may allow pied flycatchers to breed in low quality habitat and late in the season when food availability is lower (McFarlane et al 2018).

**Figure 1:**
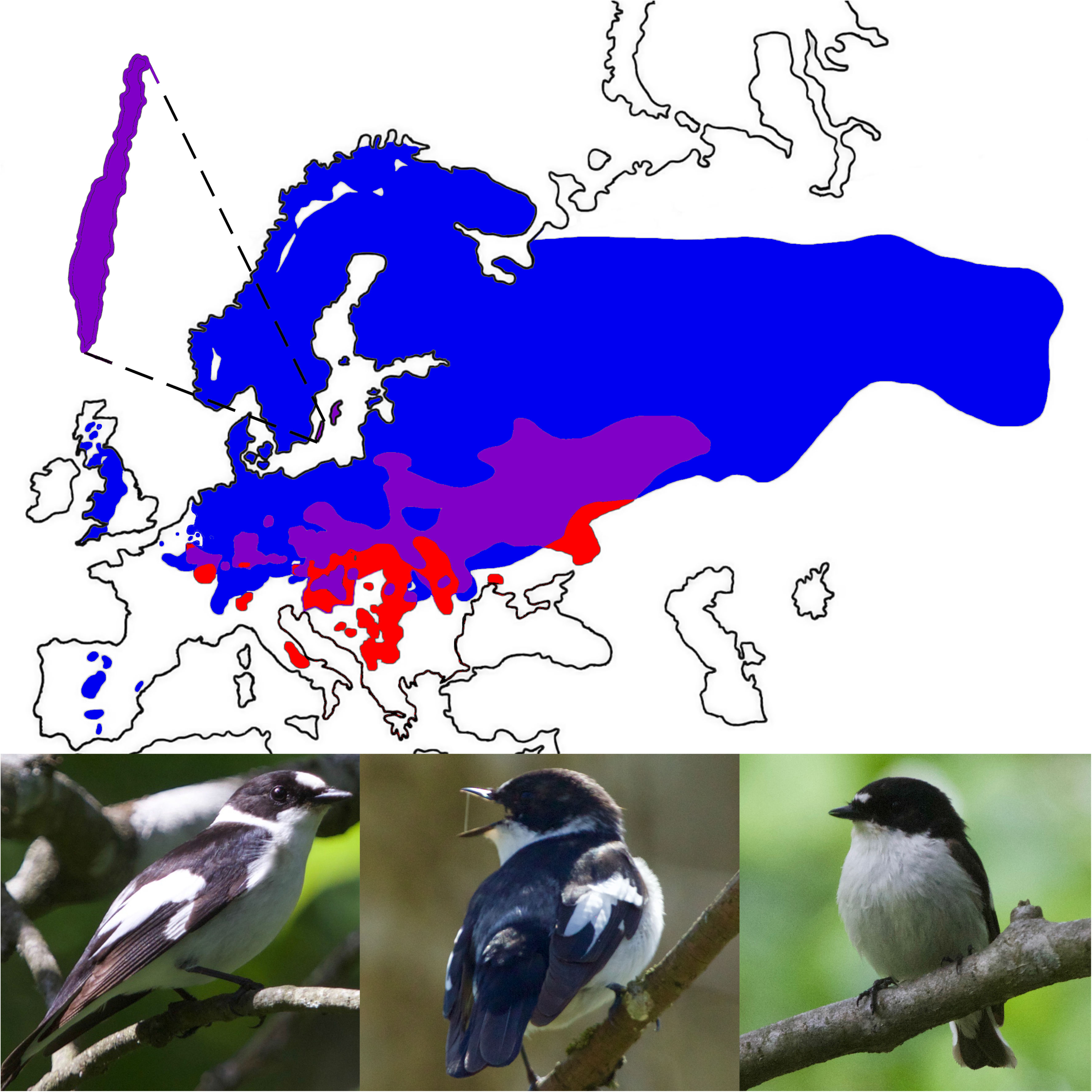
**A.** The distribution of Ficedula flycatchers in Europe. The distribution of pied flycatchers is indicated by blue, whereas the distribution of the collared flycatchers is indicated by red. The sympatric range is indicated in purple. Our study site is Öland, as enlarged in the image. **B.** A male collared flycatcher (black with a white belly, a white collar, a white forehead patch and white on its wings). **C.** A hybrid male (intermediate plumage; a broken white collar). **D.** A male pied flycatcher (black or brown with a white belly and a small white forehead patch).

Despite competition between the two species over similar nesting sites, they also interbreed at a low frequency, resulting in hybrids with an intermediate plumage phenotype (Figure 1). Hybrids display extremely low fertility as female hybrids lay empty eggs (Svedin et al 2008) and male hybrids have malformed sperm (Ålund et al 2013). Additionally, flycatcher hybrids have a higher RMR compared to both parental species (McFarlane et al 2016), suggesting increased energy requirements for maintenance and thus likely impacting fitness (i.e. less available energy for other traits). This might indicate genetic incompatibilities in the mitonuclear pathway, however, it remains unknown to what extent the two species have diverged in their mtDNA and corresponding nuclear OXPHOS genes.

In this study we aimed to investigate if genetic divergence of the mtDNA and associated nuclear OXPHOS genes between collared and pied flycatchers could play a role in hybrid dysfunction. We examined patterns of neutrality and selection on both types of DNA, to narrow in on regions that may be associated with differences in plasticity of RMR and/or be causing BDMIs between the two species.

## Materials and methods

### Population monitoring

The flycatcher population on Öland has been continuously studied since 2002 (see (Qvarnström et al 2009) for a thorough description of our field methods). Briefly, more than 2000 nestboxes are monitored for breeding activity in May and June. All birds (male and female adults, and nestlings) are individually ring marked and blood samples are taken for genetic analyses. mtDNA was sequenced from 227 individuals. Additionally, whole genome DNA data has previously been generated for 38 individuals (Burri et al 2015). All research on flycatchers was approved by the Linköping Animal Ethics Committee.

### Bioinformatics

We sequenced the 12S, 16S, ATP, CO1, CO3, CytB, ND1, ND2, ND3, ND4L, ND4 and ND6 regions of the mtDNA of 227 flycatchers (165 collared flycatchers, 45 pied flycatchers, 17 hybrids) with paired Sanger sequencing using the primers from (Amer et al 2013).

All sequences were trimmed with DNA Baser v4.36.0 (BioSoft 2013), on default settings for samples with ‘normal’ quality (trim until >60% good bases (QV>20) in 16 base window). Paired reads were assembled into contigs using the DNA Baser default settings and visually inspected for ambiguities in the forward versus the reverse strand. We obtained good quality sequences for different individuals per gene, resulting in different numbers of sequences per gene alignment.

For the nuclear analysis we used genomic sequences of 19 pied and 19 collared flycatchers from Öland (available on EMBL-EBI European Nucleotide Archive (accession PRJEB7359);(Burri et al 2015)). Exons of the OXPHOS-related genes were extracted from the genomic data aligned to the collared flycatcher reference (FicAlb1.5) using the annotated genome (we extracted both nuclear and mitochondrial genes and merged the data for the mitochondrial genes with the new sequence data). Pseudo-haploid consensus sequences of the genes were made by calling all positions covered by at least two reads and randomly choosing an allele for heterozygous sites using ANGSD–dofasta (Korneliussen et al 2014). An overview of all the included genes (nuclear and mitochondrial) can be found in Supplementary Table S1. We included all mitochondrial genes and most nuclear OXPHOS-related genes that are a part of OXPHOS complexes I, III, IV and V (complex II is not included, since this complex only contains nuclear-encoded genes and thus has no direct interactions with mitochondrial genes). One pied individual was excluded from our analysis as it clustered with collared flycatchers based on the mtDNA, suggesting potential contamination or mislabelling.

All individual contigs were aligned with AliView using MUSCLE (default settings) (Larsson 2014). The collared flycatcher mitochondrial genome (Ekblom et al 2014);(NCBI) and nuclear gene sequences from NCBI were used as reference sequences. For mtDNA, all insertions, deletions and polymorphisms in the sequences were visually inspected with DNA Baser to ensure no base calling mistakes.

### Haplotype Networks

To visualise inter- and intraspecific genetic variation we made two haplotype networks, one based on mtDNA and one based on nDNA. SequenceMatrix 1.8 (Vaidya et al 2011) was used to concatenate the genes per individual and PopART (Bandelt et al 1999, Leigh & Bryant 2015) was used to make a median-joining haplotype network of the concatenated sequences. PopART only includes sites called in >95% of individuals, so we based the mtDNA network on a concatenation of the regions for which sequences were available for most individuals (12S, ATP, ND1, ND2, ND3 and ND4) (Bandelt et al 1999, Leigh & Bryant 2015). In addition, we removed individuals that had few called bases from the analysis; the resulting haplotype network consists of 147 collared, 12 hybrid and 29 pied flycatchers. For nDNA enough sequence data was available to include all genes and individuals.

### Estimates of diversity and divergence

We calculated haplotype diversity (H_d_) and nucleotide diversity (π) as measures of genetic diversity within each species, separately for mtDNA and nuclear OXPHOS-related genes. In addition, divergence between the two species was analysed by calculating the average number of nucleotide substitutions per site between populations with a Jukes-Cantor adjustment (D_XY_) and the average number of nucleotide differences between the populations (k). All analyses were done with DnaSP (version 6.12) (Rozas et al 2017), on concatenated alignments of the genes.

mtDNA summary statistics were calculated for a concatenated alignment of the regions for which we obtained the most sequence data (12S, 16S, ATP, CO1, CO3, CytB, ND1, ND2, ND3, ND4). In order to avoid biased results due to different sample sizes we used the 23 pied and 23 collared individuals for which the most complete sequence dataset was available.

We identified and classified polymorphic sites in our alignments. A site was considered as a polymorphic site when the polymorphism was shared by either at least five individuals (when the alignment contained both newly sequenced mtDNA data and data extracted from (Burri et al 2015) (±264 individuals), or shared by at least three individuals (when an alignment only contained data extracted from (Burri et al 2015) (±37 individuals). Per site, we classified the polymorphism as a fixed difference, as variable in either species or as variable in both species, and we determined whether the site is non-coding, non-synonymous or synonymous.

Since variation in nDNA consisted for a large part of shared polymorphisms (see Results), we tested whether there was a significant difference in allele frequency for shared SNPs between the two species. Shared mitochondrial SNPs were compared between haplogroups, with mtDNA sequences for the hybrids added to the data of the corresponding maternal species (haplogroups are based on haplotype network and known species information (see Results)). We applied a chi-squared test (Ryman et al 2006) and Fisher’s exact test, which is intended for small sample sizes (Cammen et al 2015, Ryman & Jorde 2001). We used a Bonferroni correction to adjust for multiple testing. We report Fisher’s exact test below, and the chi-squared test in Supplementary Table S2. This analysis was done in R ((R Core Team 2017); scripts available in supplementary material).

### Analyses of non-neutrality and selection

Fu and Li’s F and D test statistic (FLF*, FLD*) (Fu & Li 1993) and Tajima’s D (Tajima 1989) were calculated with DnaSP to test departures from neutrality in all genes of interest. A negative neutrality test can be a sign of purifying selection, selective sweep or population expansion, whereas a positive neutrality test can be a sign of balancing selection or a bottleneck.

We used a variety of methods to identify positive selection on our genes of interest. To identify codon specific positive selection, based on the ratio of non-synonymous versus synonymous substitutions per site (ω; dN/dS), we used CodeML (from the package PAML) (Yang 2007) which uses a Bayesian Empirical Bayes-method (BEB) to identify positively selected sites. Next, we employed a likelihood ratio test to compare a model with a single ω (model 0) to models with variable ω (either nearly neutral or discrete). Subsequently, likelihood-values obtained for models including positive selection were compared to the values for models without positive selection (positive selection vs nearly neutral and beta and omega vs beta). Additionally, we used the ‘mixed effects model of evolution’ (MEME) (Murrell et al 2012) and ‘fast unconstrained Bayesian approximation’ (FUBAR) (Murrell et al 2013) from HYPHY (datamonkey.org; (Kosakovsky Pond et al 2005, Weaver et al 2018). MEME uses maximum likelihood estimation to identify episodic selection on individual sites (Murrell et al 2012). FUBAR uses a Bayesian approach to infer non-synonymous and synonymous substitution rates per site (Murrell et al 2013).

## Results

### Haplotype network

We confirmed that the two species have diverged on both the mt- and nDNA as they cluster separately in the haplotype-networks (Figures 2, 3). Based on the mtDNA network, we separated the two species into haplogroups using the pied and collared clusters, and assigned a maternal species to all hybrids. Six hybrids clustered with pied flycatchers, suggesting pied flycatcher maternity, and six hybrids clustered with collared flycatchers, suggesting collared flycatcher maternity (Figure 2). Additionally, three ‘collared hybrids’ and two ‘pied hybrids’ did not have enough sequences available for the haplotype network analysis, so are not included. Within both clusters, most haplotypes differed from each other by a few mutations, as indicated by the perpendicular bars on the edges. The collared flycatcher cluster had 17 haplotypes that were shared between several individuals, whereas in the pied flycatcher cluster most individuals had unique haplotypes (only 3 haplotypes are shared). The collared NCBI reference sequence was a part of the circle CF-10 which is the most common haplotype among collared flycatchers.

**Figure 2.**
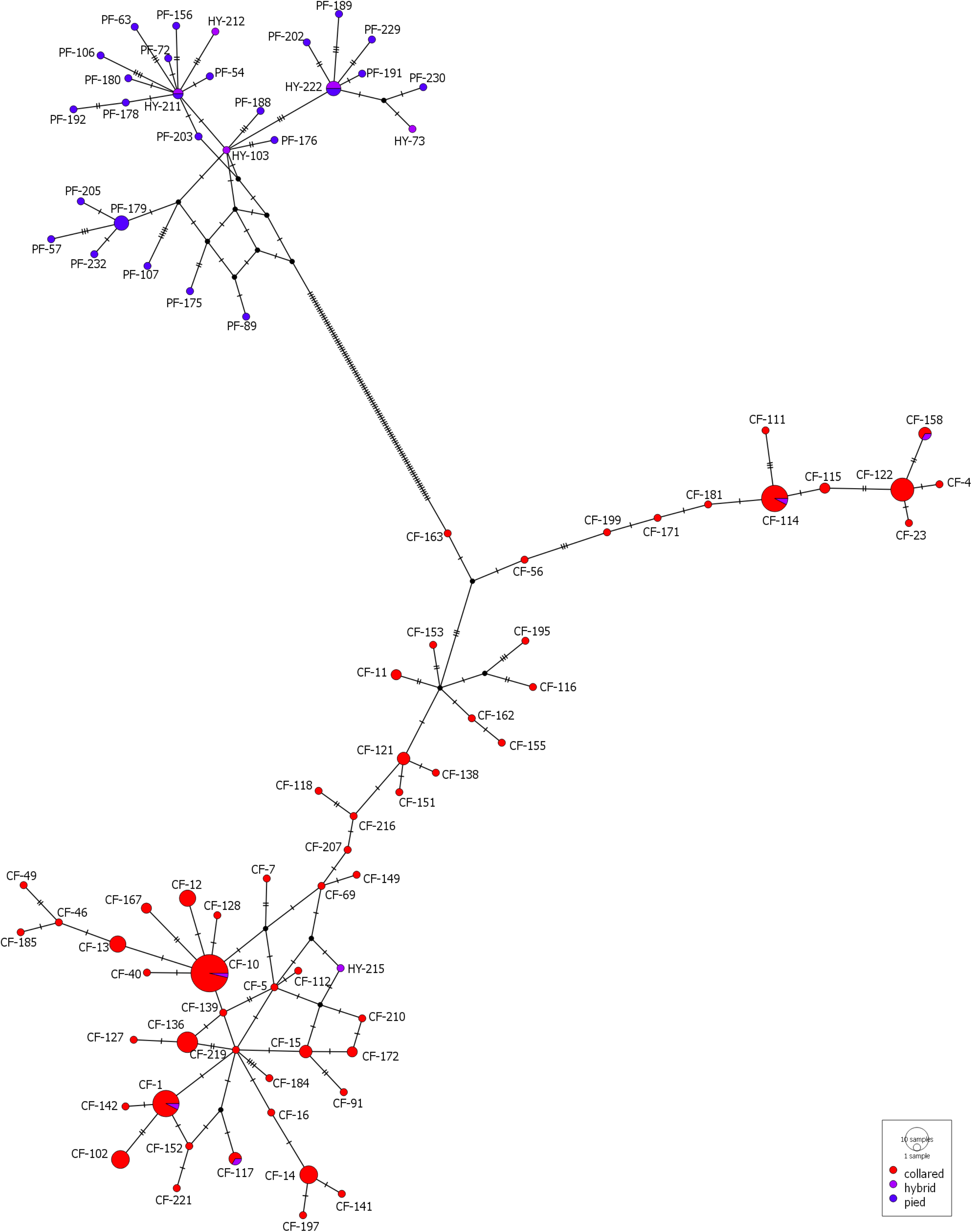
A haplotype network with all individuals based on concatenated alignment of the genes ND1, ND2, ND3, ND4, ATP and 12S. Larger circles indicate several individuals have the same haplotype. The reference sequence is present in the circle indicated by ‘CF-10’. Perpendicular bars on the edges indicate the number of differences that separates the two haplotypes. Collared and pied flycatchers cluster together based on species.

**Figure 3.**
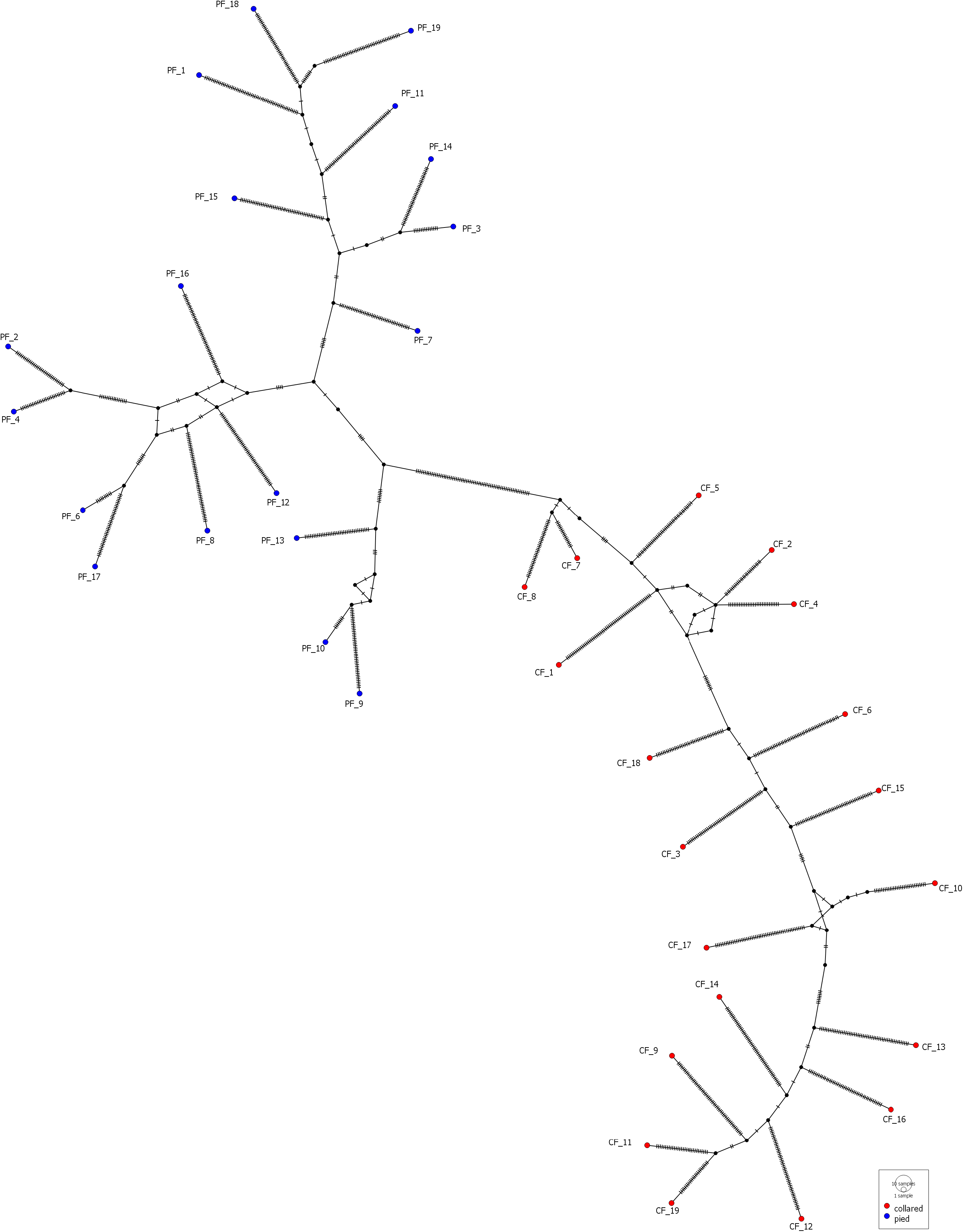
A haplotype network with all individuals based on concatenated alignment of the nuclear OXPHOS-related genes. No individuals share the same haplotype. Perpendicular bars on the edges indicate the number of differences that separates the two haplotypes. Collared and pied flycatchers cluster together based on species.

Similarly, pied and collared flycatchers cluster separately based on the nuclear haplotypes of OXPHOS-related genes (figure 3). Unsurprisingly, each individual had a unique haplotype, separated by several mutations from all other individuals.

### Genetic diversity and divergence

Mt- and nDNA showed slightly different patterns of genetic diversity and divergence. We found higher genetic and haplotype diversity in pied flycatcher mtDNA (Hd = 1, π_pied_ = 0.00162) versus collared flycatcher mtDNA (Hd = 0.976, π_collared_ = 0.00148), whereas nDNA was more diverse in collared flycatchers (π_pied_ = 0.00269, SD = 0.00008; π_collared_ = 0.00280, SD = 0.00008 (NS)) (Table 1). As expected nDNA is generally less diverged compared to mtDNA. Mitochondrial divergence between the two species measured as the absolute divergence (D_XY_) and average number of nucleotide differences between the two populations (k) (Table 1) was 0.03302 (SD = 0.00244) and 175.745 respectively. Divergence between the nDNA of the two species was 0.00422 (D_XY_, SD = 0.00031) (Table 1).

**Table 1.** Genetic diversity and divergence. Pop = gene-population; Seg = segregating sites, (st) = singletons; H = number of haplotypes; Hd = haplotype diversity; π = nucleotide diversity; D_XY_ = average number of nucleotide substitutions per site between populations (with Jukes-Cantor adjustment); k = average number of nucleotide differences between populations; (SD) = standard deviation.

| Pop | Seg (st) | H | Hd (SD) | $\pi$ (SD) | $D_{xy}$ (SD) | k |
| --- | --- | --- | --- | --- | --- | --- |
| *MT-collared* | 39 (13) | 18 | 0.976 (0.020) | 0.00148 (0.00020) | 0.03302<br>(0.00244) | 174.745 |
| *MT-pied* | 65 (55) | 22 | 1.000 (0.014) | 0.00162 (0.00015) |  |  |
| *Nuc-collared* | 311 (113) | 19 | 1.000 (0.017) | 0.00280 (0.00008) | 0.00422<br>(0.00031) | 107.125 |
| *Nuc-pied* | 254 (88) | 19 | 1.000 (0.017) | 0.00269 (0.00008) |  |  |

### Polymorphic sites

We find clear differences between the polymorphic sites in the nDNA and the mtDNA. There are fewer polymorphic sites in the nuclear OXPHOS-related genes (N = 118) compared to the mtDNA (N = 454) and more fixed differences between the species in the mtDNA (N = 188) compared to the nDNA (N = 2).

In the mtDNA, we identified 454 variable sites (out of 11,413 sites). 188 of those were fixed differences, as well as 28 shared polymorphisms and 238 polymorphisms that were monomorphic in one species but polymorphic in the other (Table S1). The majority of variable sites were in the third codon position of the nucleotide sequence and were synonymous (392/454), but we identified 62 non-synonymous polymorphisms. 13 of the non-synonymous SNPs were fixed between the two species (in ND1, ND2, ND4L, ND5, CO2 and ATP6), and a large proportion of non-synonymous polymorphisms were shared between the two species (12/28). 17 polymorphic non-synonymous mutations were polymorphic in the pied but monomorphic in the collared flycatchers, and 20 non-synonymous mutations were polymorphic in the collared but monomorphic in the pied flycatchers (Table S1).

In the coding regions of the nDNA we identified 118 variable sites (out of 20,868 sites). Many of the analysed genes did not have any variation in the coding region (Table S1). There were only two fixed differences between the two species in the coding regions of the analysed genes (both in COX7C) and five in the non-coding parts surrounding genes that were included in the alignments (ATP5F1B, ATP5MG) (Table S1). Generally, polymorphisms were shared between species (151, 46 in coding regions) although 147 (47 in coding regions) were polymorphic only in collared flycatchers and 109 (22 in coding regions) were polymorphic only in pied flycatchers. When the OXPHOS complexes were compared, eight non-synonymous mutations were present in complex I, five in complex III, no non-synonymous mutations in complex IV and one in complex V. Most non-synonymous mutations were found in collared flycatchers, with the exception of the mutation in complex V and one mutation in complex III that were only polymorphic in pied flycatchers, while five non-synonymous polymorphisms were shared between the two species.

Since many SNPs in the nDNA were shared between the two haplogroups, we calculated allele frequency differences to test for species divergence. All 179 shared SNPs (151 nDNA, 28 mtDNA) were analysed with a Fisher’s exact test and a chi-squared test (Figure 4 and Table S2). 25 SNPs were significantly different between the species, of which 15 were mtDNA and 10 nDNA. Most significant SNPs were a part of complex I (14), followed by complex IV (6), III (3) and V (2). Chi-squared tests gave very similar results, although one site in nDNA complex I was no longer significantly different (Table S2). This result indicates that even though there are fewer shared polymorphic sites in mtDNA, those sites are more often significantly different between the species.

**Figure 4.**
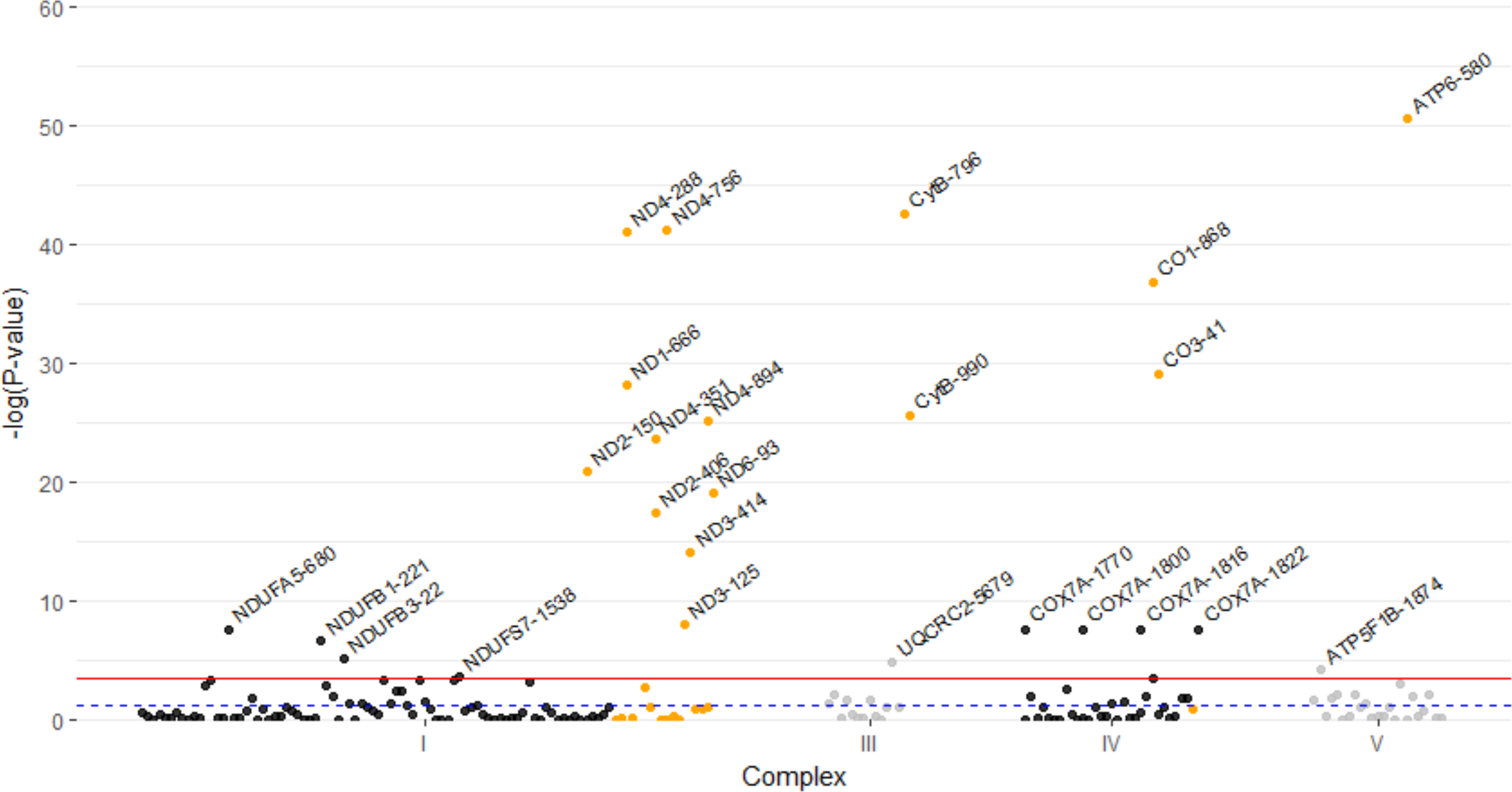
Manhattan plot with P-values of the Fisher’s Exact test for all shared SNPs. The Y-axis is the −log(P-value), and the X-axis corresponds to the complex each SNP is a part of. Black and grey SNPs are located on the nDNA, whereas orange SNPs are located on the mtDNA. The blue dashed line indicates P=0.05 and the red line indicates P=0.05/179 (P-value with Bonferroni correction), all SNPs above the red line are significantly different between the two species after accounting for multiple testing.

### Neutrality and positive selection

We found deviations from neutrality in the OXPHOS genes in the mtDNA of pied and collared flycatchers, but not in the nDNA of either species, possibly due to low power because there was little to no variation in the coding regions of the nDNA (Li’s D and F Statistics, Table 2). Specifically, in the mtDNA, we found evidence of selection on 12S, 16S, CytB, ATP6 (both species) and ND2, ND4, ND6, CO1, CO2 and CO3 (only pieds), possibly indicating a selective sweep. Alternatively, a recent population expansion might be the cause of the observed deviations from neutrality (Table 2). There was no significant pattern of selection on the other genes and the pattern was consistent between Fu and Li’s D and Tajima’s D for almost all genes (Table 2).

**Table 2.** Neutrality tests per gene and per species. FLD* = Fu and Li’s D* statistic; FLF* = Fu and Li’s F statistic; Tajima’s D = Tajima’s D neutrality statistic. Red indicates negative and significant (* = P<0.5; ** = P<0.02; *** = P<0.001), yellow is negative, green is positive, black is zero (no variation). An asterisk after the gene-name (for a mitochondrial gene) indicates only extracted data was available for that gene.

|  | Collared flycatchers |  |  | Pied flycatchers |  |  |
| --- | --- | --- | --- | --- | --- | --- |
| Gene | FLD* | FLF* | Tajima's D | FLD* | FLF* | Tajima's D |
| **Complex I** |  |  |  |  |  |  |
| **Mitochondrial genes** |  |  |  |  |  |  |
| *12S* | -3.9861** | -3.7507** | -1.6038 | -2.9181* | -3.0771** | -1.9825* |
| *16S* | -4.8867** | -4.6199** | -1.9670* | -3.2516* | -3.3062** | -1.8781* |
| *ND1* | -1.4105 | -1.4268 | -0.8288 | -1.8862 | -2.02957 | -1.3919 |
| *ND2* | -0.7600 | -0.5738 | 0.0085 | -3.2898* | -3.4395** | -2.1575* |
| *ND3* | 0.7833 | 0.8868 | 0.6837 | -1.7805 | -1.7956 | -0.9906 |
| *ND4* | -1.1539 | -1.3518 | -1.0953 | -3.1051* | -2.9786* | -1.3157 |
| *ND4L* | -2.9365* | -2.7376* | -1.1157 | -1.7957 | -2.0368 | -1.6790 |
| *ND5\** | -0.6283 | -0.5671 | -0.1460 | -0.8218 | -1.1716 | -1.5168 |
| *ND6* | -1.8019 | -1.9108 | -1.2470 | -2.6773* | -2.8955* | -2.0574* |
| **Nuclear genes** |  |  |  |  |  |  |
| *NDUFA2* | 0.0000 | 0.0000 | - | 0.0000 | 0.0000 | - |
| *NDUFA4* | -1.5203 | -1.6305 | -1.1648 | 0.0000 | 0.0000 | - |
| *NDUFA5* | -0.5736 | -0.8252 | -1.1200 | 0.0000 | 0.0000 | - |
| *NDUFA6* | 0.0000 | 0.0000 | - | 0.0000 | 0.0000 | - |
| *NDUFA8* | -1.5203 | -1.6305 | -1.1648 | 0.0000 | 0.0000 | - |
| *NDUFA11* | 0.8750 | 0.6152 | -0.3872 | 0.0000 | 0.0000 | - |
| *NDUFB1* | -0.7823 | -0.7997 | -0.4626 | 0.0094 | -0.0096 | -0.0526 |
| *NDUFB3* | -0.5736 | -0.6302 | -0.4849 | -0.1052 | -0.0231 | 0.2079 |
| *NDUFB6* | 0.0000 | 0.0000 | - | 0.0000 | 0.0000 | - |
| *NDUFB9* | -1.5203 | -1.6305 | -1.1648 | 0.0000 | 0.0000 | - |
| *NDUFB10* | 0.0000 | 0.0000 | - | 0.6578 | 0.6794 | 0.4171 |
| *NDUFS1* | 1.3036 | 1.1186 | 0.1123 | -2.2054 | -2.3859 | -1.7153 |
| *NDUFS4* | -1.2251 | -1.4688 | -1.4224 | 0.6578 | 0.6794 | 0.4171 |
| *NDUFS5* | 0.0000 | 0.0000 | - | 0.0000 | 0.0000 | - |
| *NDUFS7* | 0.0000 | 0.0000 | - | 0.0000 | 0.0000 | - |
| *NDUFS8* | 0.2317 | 0.2508 | 0.1800 | 0.0000 | 0.0000 | - |
| *NDUFV1* | 0.0000 | 0.0000 | - | 0.0000 | 0.0000 | - |
| *NDUFV2* | 0.2317 | 0.4161 | 0.6868 | -0.1432 | -0.1610 | -0.1294 |
| **Complex III** |  |  |  |  |  |  |
| **Mitochondrial genes** |  |  |  |  |  |  |
| *CytB* | -3.3245** | -3.3030** | -1.8579* | -4.2346** | -4.3038** | -2.4301** |
| **Nuclear genes** |  |  |  |  |  |  |
| *UQCRQ* | 0.0000 | 0.0000 | - | 0.0000 | 0.0000 | - |
| *UQCRC2* | -0.0279 | -0.1423 | -0.3585 | -0.7259 | -0.6050 | -0.0146 |
| *UQCR10* | 0.0000 | 0.0000 | - | 0.0000 | 0.0000 | - |
| *UQCR11* | 0.0000 | 0.0000 | - | 0.0000 | 0.0000 | - |
| **Complex IV** |  |  |  |  |  |  |
| **Mitochondrial genes** |  |  |  |  |  |  |
| *C01* | -0.9560 | -1.1432 | -0.9752 | -3.1892* | -3.1586* | -1.6174 |
| *C02\** | -0.7312 | -0.6848 | -0.2363 | -2.1748 | -2.4100 | -1.8566* |
| *C03* | -1.6433 | -1.6085 | -0.7830 | -4.1893** | -4.2057** | -2.2952** |
| **Nuclear genes** |  |  |  |  |  |  |
| *COX4I1* | -0.5736 | -0.4201 | 0.1991 | -1.5203 | -1.6305 | -1.1648 |
| *COX6C2* | 0.0000 | 0.0000 | - | 0.6578 | 0.3850 | -0.5622 |
| *COX7A* | 0.6578 | 0.3850 | -0.5622 | 0.6578 | 0.6794 | 0.4171 |
| *COX7B* | 0.0000 | 0.0000 | - | 0.0000 | 0.0000 | - |
| *COX7C* | 0.0000 | 0.0000 | - | 0.0000 | 0.0000 | - |
| **Complex V** |  |  |  |  |  |  |
| **Mitochondrial genes** |  |  |  |  |  |  |
| *ATP6* | -1.7066 | -2.1458 | -1.9573* | -4.7183** | -4.6629** | -2.3910** |
| *ATP8\** | 0.6578 | 0.6794 | 0.4171 | 0.0000 | 0.0000 | - |
| **Nuclear genes** |  |  |  |  |  |  |
| *ATP5F1B* | 0.6578 | 0.3850 | -0.5622 | 0.8750 | 1.1103 | 1.2249 |
| *ATP5F1C* | 0.8750 | 1.0353 | 0.9807 | -0.5736 | -0.4951 | -0.0452 |
| *ATP5MF* | 0.0000 | 0.0000 | - | 0.0000 | 0.0000 | - |
| *ATP5MG* | 0.0000 | 0.0000 | - | 0.0000 | 0.0000 | - |
| *ATP5PB* | 0.0000 | 0.0000 | - | 0.0000 | 0.0000 | - |
| *ATP5PD* | 0.0000 | 0.0000 | - | -2.0222 | -2.1606 | -1.5108 |
| *ATP5PF* | 0.6578 | 0.8832 | 1.0951 | 0.6578 | 0.7926 | 0.7938 |
| **Concatenated alignments** |  |  |  |  |  |  |
| *All nuclear* | -0.3591 | -0.4120 | -0.3404 | -0.2638 | -0.2290 | -0.0391 |
| *All MT* | -0.3045 | -0.4738 | -0.6294 | -3.3702** | -3.4298** | -1.9422* |

When we analysed the mtDNA genes as a concatenated alignment, FLD*, FLF* and Tajima’s D were significantly negative for the pied population, but not different from zero in the collared population (Table 2). For the concatenated nDNA OXPHOS regions, the neutrality tests for the collared flycatchers were more negative compared to the pied flycatchers (collared: FLF* = −0.3591, FLD* = −0.4120, Tajima’s D = −0.3404; pied: FLF* = −0.2638, FLD* = −0.2290, Tajima’s D = −0.0391), although none of the tests is significantly negative.

In addition to examining neutrality in each gene, we also identified positive selection on specific codons in both mtDNA and nDNA for complexes I and III. In the mitochondrial genes, we found signals of positive selection on 14 sites in ND1, ND4, CytB, CO1 and CO3 (Table 3). These genes are part of complexes I, III and IV. Further, we found positive selection between the species on 12 sites in the nuclear genes NDUFA5, NDUFB3, NDUFS1, NDUFS7, NDUFV1, UQCRC2 and ATP5PF. These genes are a part of the complexes I, III and V.

**Table 3.** Sites with positive selection. We used three programs to identify sites with positive selection, namely CodeML, MEME and FUBAR. A likelihood-ratio test (LRT) in CodeML calculated whether models with positive selection were more likely than models without positive selection. When this test was not significant, and positive selection was thus not more likely than no selection, the identified ‘positive sites’ were not accepted. In the ‘total’ column, only sites are mentioned where the LRT indicates selection and that were identified by at least two methods (the models in CodeML count as a method each). BEB = Bayesian Empirical Bayes-method. The genes for which there is no coding variation are not included in this analysis. An asterisk after the gene-name (for a mitochondrial gene) indicates only extracted data was available for that gene.

| Gene | CodeML |  |  | MEME | FUBAR | Total |
| --- | --- | --- | --- | --- | --- | --- |
|  | LRT | BEB model 2 | BEB model 8 |  |  |  |
| Complex I |  |  |  |  |  |  |
| Mitochondrial DNA |  |  |  |  |  |  |
| ND1 | Selection | 20 | 20 | 20 | 20 | 20 |
| ND2 | No selection | X | X | 30 | X | No selection |
| ND3 | No selection | X | X | X | X | No selection |
| ND4 | Selection | 4, 10, 14 | 4, 10, 14, 18 | X | 4 | 4, 10, 14 |
| ND4L | No selection | X | X | X | X | No selection |
| ND5* | Selection | 13,350,577 | 13,350,577 | 34 | X | 13,350,577 |
| ND6 | No selection | X | X | X | X | No selection |
| Nuclear DNA |  |  |  |  |  |  |
| NDUFA5 | Selection | 63 | 63 | X | X | 63 |
| NDUFB3 | Selection | 8, 57, 99 | 8, 57, 99 | X | 57 | 8, 57, 99 |
| NDUFS1 | Selection | 22, 529 | 22 | X | X | 22 |
| NDUFS5 | No selection | X | 287 | X | X | No selection |
| NDUFS7 | Selection | 21 | 21 | X | 21 | 21 |
| NDUFV1 | Selection | 7 | 7 | X | 7 | 7 |
| No coding variation: NDUFA2, NDUFA4, NDUFA6, NDUFA8, NDUFA11, NDUFB6, NDUFB9, NDUFB10, NDUFV2 |  |  |  |  |  |  |
| Complex III |  |  |  |  |  |  |
| Mitochondrial DNA |  |  |  |  |  |  |
| CytB | Selection | 23, 31, 265 | 23, 31, 265 | 265 | 31, 265 | 23, 31, 265 |
| Nuclear DNA |  |  |  |  |  |  |
| UQCRC2 | Selection | 2, 188, 311, | 2, 188, 311, 341 | 2 | 2, 188, 341 | 2, 188, 311, |
|  |  | 341 |  |  |  | 341 |
| *No coding variation: UQCRQ, UQCR10, UQCR11* |  |  |  |  |  |  |
| **Complex IV** |  |  |  |  |  |  |
| **Mitochondrial DNA** |  |  |  |  |  |  |
| *CO1* | Selection | 5, 27, 280 | 5, 27, 280 | 5 | 27 | 5, 27, 280 |
| *CO2\** | No selection | X | X | X | X | No selection |
| *CO3* | Selection | 10, 12, 18 | 10, 12, 18 | 12 | 12 | 10, 12, 18 |
| **Nuclear DNA** |  |  |  |  |  |  |
| *No coding variation: COX4I1, COX6C2, COX7A, COX7B, COX7C* |  |  |  |  |  |  |
| **Complex V** |  |  |  |  |  |  |
| **Mitochondrial DNA** |  |  |  |  |  |  |
| *ATP6* | No selection | 5, 13 | 5, 7, 13 | 13 | 13, 46 | No selection |
| *ATP8\** | No selection | X | X | X | X | No selection |
| **Nuclear DNA** |  |  |  |  |  |  |
| *ATP5PF* | Selection | 30 | 30 | X | X | 30 |
| *No coding variation: ATP5F1B, ATP5F1C, ATP5MF, ATP5MG, ATP5PB, ATP5PD* |  |  |  |  |  |  |

## Discussion

Collared and pied flycatchers diverged less than 1 million years ago (Nadachowska-Brzyska et al 2013) with strong postzygotic reproductive isolation in the form of hybrid dysfunction and infertility (Ålund et al 2013, McFarlane et al 2016, Svedin et al 2008). Here, we specifically focus on divergence in mtDNA and the interacting nuclear OXPHOS genes, as we have recently demonstrated that phenotypic differences in metabolic rate play important roles related to both differences in niche breath of the two species (McFarlane et al 2018) and hybrid dysfunction (McFarlane et al 2016). Using data from free-living collared, pied and hybrid flycatchers, we demonstrate distinct divergence in both the mtDNA (Figure 2) and in OXPHOS-related nDNA (Figure 3) of these species. Both mtDNA and nDNA divergence are significantly higher than the genome wide median divergence previously estimated for this species pair (mtDNA D_XY_ = 0.033±0.002, nDNA D_XY_ =0.0042±0.0003 compared to 0.00013 from (Ellegren et al 2012). This suggests that both the mitochondrial and nuclear OXPHOS genes have higher levels of divergence between collared and pied flycatchers than the average gene in the genome. This divergence could provide at least some of the fuel for BDMIs between the species affecting hybrid performance.

Polymorphisms with different frequencies can also contribute to incompatibilities between species (Cutter 2012). For example, a polymorphic haplotype in *Capsella sp*., which is maintained by balancing selection, results in both compatible and incompatible hybrid crosses (Sicard et al 2015). Recent work on *Drosophila* suggests that within species incompatibilities are common and are maintained at low frequencies through mutation-selection balance (Pool 2015). Incompatibilities will be kept at low frequencies within species, but could result in hybrid sterility or inviability between species if different alleles are preferred in each population. We have demonstrated here polymorphisms (both shared and not shared) in OXPHOS genes (Table S2), which could contribute to incompatibilities between collared and pied flycatchers.

While mitochondrial variation was previously thought to be rapidly purged and unlikely to influence phenotypic differences in contemporary populations, recent evidence has suggested that this variation may in fact reflect adaptive processes (Bazin et al 2006). Mitochondria are gene-dense and there is growing evidence for their non-neutral evolution (e.g. (Galtier et al 2009, Lamb et al 2018, Morales et al 2015, Ruiz-Pesini et al 2004). It is difficult to disentangle the effects of selection from the effects of demographic processes using genomic data alone without using phenotypic data to support the role of selection (Hill et al 2018, Walsh & Lynch 2018). Our study population of collared flycatchers went through a recent bottleneck as they colonized the island of Öland about fifty years ago followed by a drastic population expansion (Kardos et al 2017). In contrast, pied flycatchers were already present on the island and breed further north as compared to collared flycatchers (Figure 1). We found both higher diversity and negative values of neutrality tests in mtDNA in pied flycatchers (Table 1,2). This is not surprising given the larger starting population as well as the fact that the pied flycatcher population on Öland is not isolated to the same extent as the collared flycatcher population and hints at selection rather than demography leading to these patterns. This relatively higher diversity is consistent with the observed adaptive plasticity in pied flycatcher RMR and could be the result of a better adaption to the variable northern climate (McFarlane et al 2018). We interpret the marked divergence in OXPHOS-related genes (including fixation of different non-synonymous mutations in mtDNA) together with signs of positive selection in both mitochondrial and nuclear genes encoding OXPHOS proteins for complex I and III as a legacy of different climate adaptation from the recent time in allopatry (Qvarnström et al 2016). We found signals of selection associated with the OXPHOS complex I in both the mitochondrial and nuclear genomes (Table 2), including evidence for positive selection in ND4 and ND5 (Table 3). A similar pattern was recently found in Australian Eastern Yellow Robins (*Eopsaltria australis*), where there is a cline of mitonuclear lineages that quickly diverges (relative to dispersal distance) in a region of climatic transition, suggesting a link between mtDNA haplotype and environment in this species (Morales et al 2018). Moreover, a comprehensive meta-analysis identified ND5 as a common site of positive selection across metazoan lineages (Garvin et al 2015). Taken together, this suggests that the pattern that we have documented, especially in ND5, may indicate adaptive changes related to environmental variation.

Since the process of mitonuclear co-adaptation likely depends on climate (Dowling et al 2008), a logical prediction is that interbreeding between populations that are adapted to different climates may result in hybrid individuals with mismatched mitonuclear genes (Chou & Leu 2010). Mitonuclear mismatches have recently been hypothesized to drive reproductive isolation in the context of avian speciation processes (Hill 2017), but empirical evidence is rare. Some studies on other organisms show that poorly functioning mitonuclear interactions may lead to various forms of dysfunction ranging from increased context-dependent metabolic costs (Arnqvist et al 2010, Hoekstra et al 2013) to reduced fertility and viability of hybrids (Breeuwer & Werren 1995, Ellison & Burton 2010). These studies were all based on experimental crosses where effects of maternal mtDNA were tested against fully co-evolved versus non-coevolved nuclear background. Since introgression against a new nuclear background occurs stepwise in nature, a more likely scenario is that F1 hybrids with only partially mismatched mitonuclear genomes are already exposed to selection. This is the case with the sampled F1 *Ficedula* hybrids where mtDNA occurs together with parts of the co-evolved nuclear genome. If there could be future generations of backcrossing in flycatchers, then patterns of non-random mitonuclear co-introgression would be expected to be seen as found in Australian Eastern Yellow Robins (*Eopsaltria australis*; Morales et al 2018). These robins range across coastal and inland habitats on the southwest coast of Australia, including across variable temperature and precipitation gradients (Morales et al 2018). The underlying expectation is that backcrossed individuals with mitonuclear mismatches are continuously removed by selection. A large region of Chromosome 1A, associated with OXPHOS complex I, has co-introgressed with the associated mitolineage into the contact zone between coastal and inland populations (Morales et al 2018). This suggests that mitolineage is associated with climate adaptation in robins, a hypothesis that would be strengthened by the examination of the association between phenotypic data such as metabolic rate and these co-introgressing lineages (Hill et al 2018).

There are reported cases of metabolic dysfunction in hybrid birds, in addition to what has been reported in flycatchers. Specifically, captive stone chat hybrids (*Saxicola torquata ssp*.) with mismatched mtDNA and nuclear DNA had different RMR from the parental types (Tieleman et al 2009), and wild caught black-capped x Carolina chickadees (*Poecile atricapillus* and *P. carolinensis*) had higher mass-corrected RMR than the parental species (Olson et al 2010). When taken together, the patterns of disrupted metabolic rate in hybrid flycatchers (McFarlane et al 2016), the divergence that we have documented in the current study and hybrid sterility in *Ficedula* F1 individuals (Ålund et al 2013, Svedin et al 2008) suggest that mismatched OXPHOS interactions may have a functional effect on hybrid fitness.

Mitonuclear interactions are a likely candidate for BDMI incompatibilities across a variety of taxa (Burton and Barreto 2012), and our results are in line with this suggestion. OXPHOS genes comprise a well characterized, highly conserved gene network (Rolfe & Brown 1997, van den Heuvel & Smeitink 2001), that nonetheless has higher than expected levels of divergence between collared and pied flycatchers. This is consistent with the high level of reproductive isolation between the two flycatcher species, which have near complete postzygotic isolation after a short period of divergence. In contrast, the average hybridizing avian species pair still produces fertile hybrids until approximately 7 million years of divergence (Price & Bouvier 2002). While we have zoomed in on possible candidates for incompatibilities in this study, we have not yet found specific mitonuclear BDMIs between the two *Ficedula* species. The polymorphisms that we found, including those shared between the species, are candidates for inter-specific incompatibilities

If polymorphic incompatibilities resulting in BDMIs were driving hybrid infertility in *Ficedula* hybrids, then variation in fertility and/or RMR in the F1 generation could be associated with variation among mismatched mitonuclear genotypes to pinpoint specific BDMIs. Additionally, we might expect to find some fertile hybrids, as has been documented both in the above *Capsella* hybrids as well as in *Mus musculus musculus* × *M. m. domesticus* where polymorphic incompatibilities have also been documented (Larson et al 2018). However, all sampled flycatcher hybrids appear to be infertile (Ålund et al 2013, Svedin et al 2008) suggesting that if there are compatible haplotypes at these loci, they are rare. In order to specify potential specific mitonuclear BDMIs caused by fixed differences between the two *Ficedula* species, we would need to use genomic methods that could restore hybrid fitness such as laboratory methods that would allow us to have two copies of candidate alleles against an otherwise F1 background. This is, however, beyond the scope of the current study.

At late stages of speciation (i.e. when postzygotic isolation is complete or nearly complete), it is difficult to tell whether specific incompatibilities have evolved before or after reproductive isolation (Seehausen et al 2014), as BDMIs tend to ‘snowball’. Theoretically, it may only take one epistatic interaction stemming from divergence while in allopatry to lead to complete reproductive isolation (Bateson 1909, Dobzhansky 1936, Muller 1940, 1942). After isolation is complete, other incompatibilities can accumulate, but these are a consequence rather than a cause of reproductive isolation, leading to the build-up of tens or hundreds of genes involved in incompatibilities (Presgraves 2003). For example, hundreds of genes affect hybrid incompatibilities between recently diverged swordtail species (*Xiphophorus birchmanni* and *X. malinche*; Schumer et al 2014). While it seems possible that the divergence in OXPHOS genes is due to differences in climate adaptation between the flycatcher species (Qvarnström et al 2016), it is premature to conclude whether mitonuclear incompatibilities are causing postzygotic isolation between the two species.

Here, we demonstrate higher than expected divergence between collared and pied flycatchers in the OXPHOS genes, hinting that these genes may be contributing to the hybrid metabolic dysfunction previously documented, possibly via BDMIs. We also found evidence of recent selection, particularly in the mtDNA gene ND5 which is part of the OXPHOS complex I and appears to be diverging between collared and pied flycatchers, consistent with selection found in other avian studies. Taken together, this suggests that metabolic dysfunction resulting from mt-nDNA incompatibilities may be a factor contributing to hybrid dysfunction in this system.

## Author Contributions

SEM and AQ conceived and designed the study and collected the data. Evdh and TvdV performed analyses. Evdh, SEM and AQ wrote the manuscript and all authors contributed to meaningful discussions and revisions of the manuscript.

## Supporting information

allele frequency analyses in R

## Acknowledgements

We are thankful to all flycatcher field assistants who contributed to the collection of these data, to Reija Dufva who conducted the molecular laboratory work, and Carolina Segami for helpful discussions. This work was funded by Vetenskapsrådet to AQ: DR 2016 −05138 and SEM: International Postdoc 2017-00499.

## Abbreviations

mtDNA: mitochondrial DNA
nDNA: nuclear DNA
BDMI: Bateson Dobzhansky Muller incompatibilities
RMR: resting metabolic rate
OXPHOS: oxidative phosphorylation pathway
FLF*: Fu’s and Li’s F* statistic
FLD*: Fu’s and Li’s D* statistic
SNP: single nucleotide polymorphism

