## Supplementary material for "Divergent mitochondrial and nuclear OXPHOS genes are candidates for genetic incompatibilities in *Ficedula* Flycatchers": allele frequency analyses in R

### R scripts for van der heijden et al. 
### these scripts perform fisher's exact tests with corresponding manhattan plot and chi squared tests. 

#libraries
library(readxl)
library("qqman", lib.loc="~/R/win-library/3.3")
library(ggplot2)
library(dplyr)


#import data --> one file for collared, one for pied 
collared_sharedSNPs <- read_excel("C:/Users/vande/OneDrive/Documenten/School/MEME/Uppsala 2/shared/all shared SNPs.xlsx", sheet = "collared")
pied_sharedSNPs <- read_excel("C:/Users/vande/OneDrive/Documenten/School/MEME/Uppsala 2/shared/all shared SNPs.xlsx", sheet = "pied")

#every column is one SNP, delete first column with ID; set all missing data to NA 
collared_sharedSNPs[,1] <- NULL
pied_sharedSNPs[,1] <- NULL
collared_sharedSNPs[collared_sharedSNPs == "N"] <- NA
pied_sharedSNPs[pied_sharedSNPs == "N"] <- NA

#import SNP names (all SNPs, all mitochondrial SNPs), will be used for graph
SNP <- read_excel("C:/Users/vande/OneDrive/Documenten/School/MEME/Uppsala 2/shared/all shared SNPs.xlsx", sheet = "SNPs")
MT_SNPs <- read_excel("C:/Users/vande/OneDrive/Documenten/School/MEME/Uppsala 2/shared/all shared SNPs.xlsx", sheet = "MT-SNPs")
MT_SNP <- as.vector(t(MT_SNPs), mode = "character")

#calculate number of SNPs, this is usefull as a variable
N_SNPs <- length(t(SNP))

### Perform Fisher's exact test for all SNPs and store the p-values
p_vals <- c()
for (i in 1:N_SNPs) {
  result <-  fisher.test(cbind(table(collared_sharedSNPs[,i]),table(pied_sharedSNPs[,i])))
  p_vals <- c(p_vals, result$p.value)
}

### Make Manhattan plot with ggplot
#make dataframe with necessary information from P-values
#NOTE: BP is just a number, not an actual base position, CHR is the complex, not chromosome!
#I have been playing around with BP to make it look nice in the graph
p_values <- as.data.frame(p_vals)
p_values$BP <- c(c(1:86,78,88:98,85,99,90,92,99,100),          #1:100
                 c(120:134),                                   #120:134
                 c(154:167,seq(154,184,10),168:183,176,177),   #154:184
                 c(204:226,220))                               #204:226
p_values$CHR <- c(rep(1,104),rep(3,15),rep(4,36),rep(5,24))
p_values <- cbind(p_values, SNP)

#Add a column with significant SNPs, to give them a label in the graph
sign <- c()
for (i in 1:N_SNPs) {
  result <-  if(p_values$p_vals[i]<=0.05/N_SNPs) as.character(p_values$SNP[i]) else ""
  sign <- c(sign, result)
}

p_values$sign <- sign

#Highlight mtDNA SNPs
p_values$highlight <- ifelse(p_values$SNP %in% MT_SNPs$MTSNPs, "yes", "no")
subset_mt <- subset(x=p_values, subset=highlight=="yes")

CENTER <- c(50,127,169,215)
CHR <- c("I","III","IV","V")

ggplot(p_values, aes(x=BP, y=-log10(p_vals), label = p_values$sign)) +
  xlab("Complex") +
  ylab("-log(P-value)")  +
  geom_point(aes(color=as.factor(CHR)), alpha=0.8, size=1.3) +
  geom_point(data=subset_mt,
             aes(x=subset_mt$BP, y=-log10(subset_mt$p_vals),
             label=subset_mt$sign), colour="orange") +
  scale_color_manual(values = rep(c("black", "grey"), 3)) +
  scale_x_continuous(label=CHR, breaks=CENTER) +
  scale_y_continuous(expand=c(0,0), limits=c(0,60)) +
  geom_hline(yintercept =-log10(0.05), col="blue", linetype="dashed") +
  geom_hline(yintercept = -log10(0.05/N_SNPs), col="red") +
  theme_bw() +
  theme( 
    legend.position="none",
    panel.border = element_blank(),
    panel.grid.major.x = element_blank(),
    panel.grid.minor.x = element_blank()
  ) +
  geom_text(hjust=0, nudge_x=1, nudge_y=1, angle=35, size=3)
  

##CHI-SQUARE TEST
### Perform chi-squared test and store the p-values
p_vals_chisq <- c()
for (i in 1:N_SNPs) {
  result_chisq <-  chisq.test(cbind(table(collared_sharedSNPs[,i]),table(pied_sharedSNPs[,i])))
  p_vals_chisq <- c(p_vals_chisq, result_chisq$p.value)
}

#Add column whether SNP is significant or not, for fisher and chisquare data
sign.fisher <- c()
for (i in 1:N_SNPs) {
  result <-  if(p_vals[i]<=0.05/N_SNPs) "*" else ""
  sign.fisher <- c(sign.fisher, result)
}

sign.chisq <- c()
for (i in 1:N_SNPs) {
  result <-  if(p_vals_chisq[i]<=0.05/N_SNPs) "*" else ""
  sign.chisq <- c(sign.chisq, result)
}

#Make a table with Fisher results and Chisquare results
table_allele_freq <- SNP
table_allele_freq$fisher <- p_vals
table_allele_freq$sign.fisher <- sign.fisher
table_allele_freq$chisq <- p_vals_chisq2
table_allele_freq$sign.chisq <- sign.chisq

write.table(table_allele_freq, "~/allele_freq.txt", sep="\t")
