## Supplementary Information and Tables for "Divergent mitochondrial and nuclear OXPHOS genes are candidates for genetic incompatibilities in *Ficedula* Flycatchers"

**Supplementary material**

Script 1 – Fisher’s exact tests and chi-squared tests

We did a Fisher’s exact test for 179 shared SNPs. We used the package ‘ggplot2’ to make a Manhattan-plot to show the P-values on a negative log-scale. We did a Bonferroni-adjustment by setting the ‘genome-wide significant’ line to 0.05/179. All SNPs with a -log(P-value) above -log(0.05/179) will be considered significant.

A chi-squared analysis was also performed on the 179 shared SNPs. The P-values for both Fisher’s exact and chi-squared analyses were combined into a table for supplementary materials.

| **Table S1. Types of polymorphism per gene**  For every gene we looked at the type of polymorphisms that were present. The four categories polymorphisms were divided in are ‘fixed differences’ (monomorphic in both species but for different bases), ‘collared poly – pied mono’ (polymorphic for collared individuals and monomorphic for pied individuals), ‘collared mono – pied poly’ (monomorphic for collared and polymorphic for pieds), and ‘shared mutations’ (polymorphic in both species). Per category, the first number refers to the number of synonymous mutations in that category, the second number refers to the number of non-synonymous mutations and the third number refers to the number of non-coding mutations.  An asterisk after the gene-name (for a mitochondrial gene) indicates only extracted data was available for that gene. | | | | | | |
| --- | --- | --- | --- | --- | --- | --- |
| **Gene** | **NCBI gene ID** | **Fixed difference** | **Collared poly – pied mono** | **Collared mono – pied poly** | **Shared variation** | **Total** |
| **Complex I** | | | | | | |
| **Mitochondrial DNA** | | | | | | |
| *ND1* | 16027862 | 16 / 1 / 0 | 8 / 0 / 0 | 11 / 1 / 0 | 2 / 2 / 0 | 37 / 4 / 0 |
| *ND2* | 16027866 | 22 / 4 / 0 | 11 / 2 / 0 | 16 / 2 / 0 | 3 / 5 / 0 | 52 / 13 / 0 |
| *ND3* | 16027881 | 4 / 0 / 0 | 3 / 0 / 0 | 5 / 2 / 0 | 1 / 0 / 1 | 13 / 2 / 1 |
| *ND4* | 16027884 | 16 / 0 / 0 | 8 / 6 / 0 | 13 / 2 / 0 | 3 / 3 / 0 | 40 / 11 / 0 |
| *ND4L* | 16027883 | 2 / 1 / 0 | 2 / 0 / 0 | 7 / 1 / 0 | 1 / 0 / 0 | 12 / 2 / 0 |
| *ND5** | 16027888 | 35 / 4 / 0 | 5 / 5 / 0 | 26 / 3 / 0 | 0 / 0 / 0 | 66 / 12 / 0 |
| *ND6* | 16027892 | 4 / 0 / 0 | 3 / 0 / 0 | 6 / 1 / 0 | 1 / 0 / 0 | 14 / 1 / 0 |
| **Nuclear DNA** | | | | | | |
| *NDUFA2* | 101811084 | 0 / 0 / 0 | 0 / 0 / 0 | 0 / 0 / 0 | 0 / 0 / 0 | 0 / 0 / 0 |
| *NDUFA4* | 101807875 | 0 / 0 / 0 | 0 / 0 / 4 | 0 / 0 / 6 | 0 / 0 / 9 | 0 / 0 / 19 |
| *NDUFA5* | 101818568 | 0 / 0 / 0 | 0 / 1 / 4 | 0 / 0 / 5 | 0 / 0 / 5 | 0 / 1 / 14 |
| *NDUFA6* | 101820913 | 0 / 0 / 0 | 0 / 0 / 2 | 1 / 0 / 0 | 1 / 0 / 2 | 2 / 0 / 4 |
| *NDUFA8* | 101821747 | 0 / 0 / 0 | 0 / 0 / 0 | 0 / 0 / 0 | 0 / 0 / 1 | 0 / 0 / 1 |
| *NDUFA11* | 101816329 | 0 / 0 / 0 | 1 / 0 / 0 | 0 / 0 / 0 | 0 / 0 / 0 | 1 / 0 / 0 |
| *NDUFB1* | 101822125 | 0 / 0 / 0 | 3? / 3? / 4 | 0 / 0 / 3 | 0 / 0 / 15 | 3? / 3? / 22 |
| *NDUFB3** | 101816515 | 0 / 0 / 0 | 0 / 1 / 0 | 0 / 0 / 0 | 0 / 2 / 0 | 0 / 3 / 0 |
| *NDUFB6** | 101809750 | 0 / 0 / 0 | 0 / 0 / 0 | 0 / 0 / 0 | 0 / 0 / 2 | 0 / 0 / 2 |
| *NDUFB9** | 101821515 | 0 / 0 / 0 | 0 / 0 / 2 | 0 / 0 / 5 | 1 / 0 / 7 | 1 / 0 / 15 |
| *NDUFB10** | 101806897 | 0 / 0 / 0 | 2 / 0 / 0 | 0 / 0 / 0 | 2 / 0 / 0 | 4 / 0 / 0 |
| *NDUFS1** | 101808800 | 0 / 0 / 0 | 3 / 1 / 4 | 1 / 0 / 7 | 0 / 0 / 3 | 4 / 1 / 14 |
| *NDUFS4** | 101819987 | 1? / 1? / 0 | 0 / 0 / 0 | 1? / 1? / 0 | 0 / 0 / 0 | 2? / 2? / 0 |
| *NDUFS5** | 101813310 | 0 / 0 / 0 | 3 / 1 / 1 | 3 / 0 / 1 | 3 / 0 / 1 | 9 / 1 / 3 |
| *NDUFS7** | 101819119 | 0 / 0 / 0 | 1 / 1 / 10 | 0 / 0 / 7 | 0 / 0 / 11 | 1 / 1 / 28 |
| *NDUFS8** | 101808792 | 0 / 0 / 0 | 0 / 0 / 8 | 0 / 0 / 0 | 2? / 2? / 4 | 2? / 2? / 12 |
| *NDUFV1** | 101812104 | 0 / 0 / 0 | 2 / 0 / 1 | 2 / 0 / 1 | 4 / 1 / 2 | 8 / 1 / 4 |
| *NDUFV2** | 101811184 | 0 / 0 / 0 | 2 / 0 / 0 | 1 / 0 / 0 | 4 / 0 / 0 | 7 / 0 / 0 |
| **Complex III** | | | | | | |
| **Mitochondrial DNA** | | | | | | |
| *CytB* | 16027889 | 21 / 0 / 0 | 6 / 3 / 0 | 12 / 2 / 0 | 1 / 1 / 0 | 40 / 6 / 0 |
| **Nuclear DNA** | | | | | | |
| *UQCRQ** | 101815283 | 0 / 0 / 0 | 0 / 0 / 0 | 0 / 0 / 0 | 0 / 0 / 0 | 0 / 0 / 0 |
| *UQCRC2** | 101822003 | 0 / 0 / 0 | 3 / 1 / 25 | 1 / 2 / 26 | 2 / 2 / 9 | 6 / 5 / 60 |
| *UQCR10** | 101817945 | 0 / 0 / 0 | 1 / 0 / 0 | 1 / 0 / 0 | 0 / 0 / 0 | 2 / 0 / 0 |
| *UQCR11** | 101805950 | 0 / 0 / 0 | 1 / 0 / 0 | 0 / 0 / 0 | 0 / 0 / 0 | 1 / 0 / 0 |
| **Complex IV** | | | | | | |
| **Mitochondrial DNA** | | | | | | |
| *CO1* | 16027872 | 25 / 0 / 0 | 6 / 0 / 0 | 13 / 0 / 0 | 1 / 1 / 0 | 45 / 1 / 0 |
| *CO2** | 16027875 | 10 / 1 / 0 | 3 / 0 / 0 | 3 / 0 / 0 | 0 / 0 / 0 | 16 / 1 / 0 |
| *CO3* | 16027879 | 12 / 0 / 0 | 2 / 4 / 0 | 10 / 0 / 0 | 1 / 0 / 0 | 25 / 4 / 0 |
| **Nuclear DNA** | | | | | | |
| *COX4I1** | 101822066 | 0 / 0 / 0 | 1 / 0 / 3 | 0 / 0 / 3 | 1 / 0 / 7 | 2 / 0 / 13 |
| *COX6C2** | 101815193 | 0 / 0 / 0 | 0 / 0 / 1 | 0 / 0 / 0 | 0 / 0 / 3 | 0 / 0 / 4 |
| *COX7A** | 101809314 | 0 / 0 / 0 | 0 / 0 / 5 | 1 / 0 / 10 | 0 / 0 / 10 | 1 / 0 / 25 |
| *COX7B** | 101811767 | 0 / 0 / 0 | 0 / 0 / 6 | 0 / 0 / 3 | 0 / 0 / 12 | 0 / 0 / 21 |
| *COX7C** | 101807117 | 2 / 0 / 0 | 0 / 0 / 2 | 0 / 0 / 1 | 0 / 0 / 0 | 2 / 0 / 3 |
| **Complex V** | | | | | | |
| **Mitochondrial DNA** | | | | | | |
| *ATP6* | 16027878 | 8 / 2 / 0 | 1 / 0 / 0 | 16 / 1 / 0 | 1 / 0 / 0 | 26 / 3 / 0 |
| *ATP8** | 16027877 | 0 / 0 / 0 | 1 / 0 / 0 | 4 / 2 / 0 | 0 / 0 / 0 | 5 / 2 / 0 |
| **Nuclear DNA** | | | | | | |
| *ATP5F1B** | 101820650 | 0 / 0 / 2 | 1 / 0 / 18 | 2 / 0 / 8 | 0 / 0 / 7 | 3 / 0 / 35 |
| *ATP5F1C** | 101808452 | 0 / 0 / 0 | 1 / 0 / 9 | 1 / 0 / 2 | 1 / 0 / 13 | 3 / 0 / 24 |
| *ATP5MF** | 101810684 | 0 / 0 / 0 | 0 / 0 / 0 | 2 / 0 / 0 | 0 / 0 / 0 | 2 / 0 / 0 |
| *ATP5MG** | 101812199 | 0 / 0 / 3 | 1 / 0 / 0 | 0 / 0 / 1 | 0 / 0 / 0 | 1 / 0 / 4 |
| *ATP5PB** | 101816794 | 0 / 0 / 0 | 2 / 0 / 0 | 0 / 0 / 0 | 1 / 0 / 0 | 3 / 0 / 0 |
| *ATP5PD** | 101815229 | 0 / 0 / 0 | 0 / 0 / 3 | 0 / 0 / 1 | 0 / 0 / 1 | 0 / 0 / 5 |
| *ATP5PF** | 101812412 | 0 / 0 / 0 | 1 / 0 / 0 | 0 / 1 / 0 | 0 / 0 / 0 | 1 / 1 / 0 |

| **Table S2. Results Fisher exact and Chi-Square for allele frequency**  The P-values per shared SNP for both the Fisher Exact and the Chi-Square test. These tests were done to test whether there is a significant difference in allele frequency for shared SNPs between the two species. An asterisk (*) indicates the test was significant after a Bonferroni-adjustment. | | | | | | | |
| --- | --- | --- | --- | --- | --- | --- | --- |
|  | **SNP** | **Fisher** | **Chi-square** |  | **SNP** | **Fisher** | **Chi-square** |
| 1 | ATP6-580 | 2.78E-51* | 1.55E-49* | 50 | COX7B-1473 | 0.01522 | 0.029413 |
| 2 | CytB-796 | 2.67E-43* | 3.17E-43* | 51 | UQCRC2-4524 | 0.01704 | 0.028669 |
| 3 | ND4-756 | 6.20E-42* | 1.08E-38* | 52 | ATP5F1B-1024 | 0.018752 | 0.037172 |
| 4 | ND4-288 | 9.84E-42* | 3.81E-34* | 53 | UQCRC2-665 | 0.02171 | 0.032097 |
| 5 | CO1-868 | 1.40E-37* | 4.20E-41* | 54 | COX7B-5 | 0.027491 | 0.02994 |
| 6 | CO3-41 | 9.43E-30* | 7.03E-32* | 55 | NDUFS1-3554 | 0.029298 | 0.040179 |
| 7 | ND1-666 | 6.84E-29* | 1.37E-24* | 56 | NDUFB6-477 | 0.034215 | 0.033564 |
| 8 | CytB-990 | 2.59E-26* | 4.61E-29* | 57 | NDUFB9-1029 | 0.038199 | 0.051663 |
| 9 | ND4-894 | 6.26E-26* | 1.29E-22* | 58 | COX7A-2030 | 0.042245 | 0.05599 |
| 10 | ND4-351 | 1.94E-24* | 1.69E-21* | 59 | NDUFB3-169 | 0.04371 | 0.045167 |
| 11 | ND2-150 | 1.07E-21* | 4.39E-20* | 60 | UQCRC2-6 | 0.044812 | 0.068179 |
| 12 | ND6-93 | 7.32E-20* | 2.77E-19* | 61 | ATP5F1C-1131 | 0.044812 | 0.060929 |
| 13 | ND2-406 | 4.09E-18* | 6.37E-20* | 62 | NDUFS7-1618 | 0.060452 | 0.094558 |
| 14 | ND3-414 | 7.68E-15* | 2.26E-15* | 63 | NDUFB9-1139 | 0.062496 | 0.079857 |
| 15 | ND3-125 | 9.74E-09* | 8.09E-10* | 64 | NDUFB9-301 | 0.067112 | 0.101953 |
| 16 | NDUFA5-680 | 2.05E-08* | 3.48E-07* | 65 | UQCRC2-5764 | 0.075036 | 0.116063 |
| 17 | COX7A-1770 | 2.05E-08* | 3.48E-07* | 66 | NDUFS7-1615 | 0.075868 | 0.119965 |
| 18 | COX7A-1800 | 2.05E-08* | 3.48E-07* | 67 | COX7B-485 | 0.076812 | 0.100198 |
| 19 | COX7A-1816 | 2.05E-08* | 3.48E-07* | 68 | NDUFV2-531 | 0.078891 | 0.100457 |
| 20 | COX7A-1822 | 2.05E-08* | 3.48E-07* | 69 | ATP5F1C-1468 | 0.078891 | 0.122051 |
| 21 | NDUFB1-221 | 1.95E-07* | 1.79E-06* | 70 | COX7A-35 | 0.083333 | 0.160581 |
| 22 | NDUFB3-22 | 6.71E-06* | 3.25E-05* | 71 | COX4I1-385 | 0.084008 | 0.140734 |
| 23 | UQCRC2-5679 | 1.26E-05* | 5.39E-05* | 72 | ND2-331 | 0.085407 | 0.133871 |
| 24 | ATP5F1B-1874 | 5.48E-05* | 0.000157* | 73 | ND4-298 | 0.085947 | 0.159527 |
| 25 | NDUFS7-1538 | 0.000234* | 0.000541 | 74 | NDUFS8-1238 | 0.086998 | 0.147892 |
| 26 | COX7B-278 | 0.000297 | 0.000922 | 75 | NDUFB1-54 | 0.088621 | 0.158259 |
| 27 | NDUFA5-573 | 0.000364 | 0.001054 | 76 | UQCRC2-5481 | 0.091098 | 0.133869 |
| 28 | NDUFS1-2559 | 0.000364 | 0.001054 | 77 | ATP5F1C-1130 | 0.091098 | 0.117067 |
| 29 | NDUFS7-1073 | 0.000364 | 0.001054 | 78 | ND4L-288 | 0.101414 | 0.195508 |
| 30 | NDUFB9-994 | 0.000451 | 0.000846 | 79 | ND4-4 | 0.101414 | 0.195508 |
| 31 | NDUFS8-358 | 0.000611 | 0.001286 | 80 | NDUFS5-1137 | 0.104927 | 0.169099 |
| 32 | ATP5F1C-1990 | 0.00083 | 0.001741 | 81 | CO1-170 | 0.111883 | 0.142107 |
| 33 | NDUFA5-551 | 0.001098 | 0.002831 | 82 | NDUFB1-9 | 0.120588 | 0.233632 |
| 34 | NDUFB1-270 | 0.001098 | 0.002831 | 83 | NDUFS7-1614 | 0.14107 | 0.175536 |
| 35 | ND2-324 | 0.00179 | 0.00129 | 84 | ATP5F1C-2246 | 0.146353 | 0.179417 |
| 36 | COX4I1-565 | 0.002398 | 0.00336 | 85 | NDUFB9-625 | 0.151026 | 0.182768 |
| 37 | NDUFB9-1030 | 0.002918 | 0.005244 | 86 | NDUFA6-1069 | 0.170447 | 0.242851 |
| 38 | NDUFB9-1035 | 0.002918 | 0.005244 | 87 | NDUFB1-55 | 0.179025 | 0.288227 |
| 39 | ATP5F1B-3767 | 0.00671 | 0.007567 | 88 | NDUFS8-270 | 0.1874 | 0.26316 |
| 40 | ATP5F1C-2324 | 0.00671 | 0.011363 | 89 | NDUFA4-908 | 0.191355 | 0.242648 |
| 41 | UQCRC2-476 | 0.006941 | 0.012348 | 90 | COX7B-213 | 0.224503 | 0.262438 |
| 42 | ATP5F1C-713 | 0.007816 | 0.012697 | 91 | NDUFA4-617 | 0.231989 | 0.357965 |
| 43 | ATP5F1C-2022 | 0.008138 | 0.013614 | 92 | NDUFV1-204 | 0.235165 | 0.28343 |
| 44 | NDUFB1-1541 | 0.010432 | 0.014047 | 93 | NDUFV2-412 | 0.269065 | 0.389858 |
| 45 | COX4I1-96 | 0.010432 | 0.014047 | 94 | NDUFB1-60 | 0.280014 | 0.365089 |
| 46 | COX7B-231 | 0.011261 | 0.010509 | 95 | NDUFS1-211 | 0.288988 | 0.288844 |
| 47 | NDUFA8-779 | 0.012767 | 0.023461 | 96 | UQCRC2-1874 | 0.295296 | 0.408466 |
| 48 | ATP5F1B-2632 | 0.013954 | 0.019484 | 97 | NDUFS7-2080 | 0.300699 | 0.468717 |
| 49 | COX7B-1472 | 0.01522 | 0.029413 | 98 | COX7B-332 | 0.313725 | 0.444359 |
|  | **SNP** | **Fisher** | **Chisquare** |  | **SNP** | **Fisher** | **Chisquare** |
| 99 | NDUFB9-777 | 0.324544 | 0.420397 | 140 | NDUFA4-910 | 0.728106 | 0.812365 |
| 100 | NDUFA4-806 | 0.339768 | 0.557279 | 141 | NDUFA6-506 | 0.728106 | 0.914812 |
| 101 | COX6C2-36 | 0.374684 | 0.483666 | 142 | COX6C2-141 | 0.730014 | 0.980492 |
| 102 | NDUFB1-43 | 0.377709 | 0.442063 | 143 | ND1-825 | 0.735697 | 0.761847 |
| 103 | ATP5F1B-5077 | 0.389847 | 0.445102 | 144 | ND1-169 | 0.735924 | 0.69159 |
| 104 | NDUFB1-49 | 0.400323 | 0.491478 | 145 | NDUFA4-906 | 0.743136 | 0.850554 |
| 105 | NDUFB10-142 | 0.417284 | 0.612946 | 146 | NDUFA4-905 | 0.745855 | 0.879484 |
| 106 | NDUFA4-621 | 0.447032 | 0.627054 | 147 | UQCRC2-2498 | 0.745855 | 0.879484 |
| 107 | ATP5F1C-1364 | 0.447032 | 0.627054 | 148 | UQCRC2-5193 | 1 | 0.995295 |
| 108 | ND2-651 | 0.451919 | 0.653506 | 149 | NDUFA4-794 | 1 | 1 |
| 109 | NDUFV1-1352 | 0.454273 | 0.66619 | 150 | NDUFA4-912 | 1 | 1 |
| 110 | COX7B-929 | 0.475634 | 0.540105 | 151 | NDUFB1-3 | 1 | 1 |
| 111 | ATP5F1C-1464 | 0.475634 | 0.540105 | 152 | NDUFB1-35 | 1 | 1 |
| 112 | ATP5F1C-2138 | 0.495055 | 0.641747 | 153 | NDUFB1-61 | 1 | 1 |
| 113 | ATP5F1B-2431 | 0.505085 | 0.620864 | 154 | NDUFB1-69 | 1 | 1 |
| 114 | COX7A-1690 | 0.505495 | 1 | 155 | NDUFB1-1571 | 1 | 1 |
| 115 | COX7A-2015 | 0.514813 | 0.63442 | 156 | NDUFB6-23 | 1 | 1 |
| 116 | NDUFV2-57 | 0.517124 | 0.624763 | 157 | NDUFS5-1353 | 1 | 1 |
| 117 | UQCRC2-5120 | 0.517124 | 0.624763 | 158 | NDUFS5-1848 | 1 | 1 |
| 118 | NDUFV1-752 | 0.544156 | 0.730385 | 159 | NDUFS5-2530 | 1 | 1 |
| 119 | ATP5PD-35 | 0.545455 | 0.540291 | 160 | NDUFS7-2084 | 1 | 1 |
| 120 | NDUFS7-2228 | 0.555769 | 0.77432 | 161 | NDUFS7-2356 | 1 | 1 |
| 121 | NDUFB1-76 | 0.564532 | 0.805972 | 162 | NDUFS8-1227 | 1 | 0.965724 |
| 122 | ATP5PB-584 | 0.601533 | 0.693034 | 163 | NDUFV1-240 | 1 | 1 |
| 123 | NDUFA5-595 | 0.603861 | 0.961038 | 164 | NDUFV1-1244 | 1 | 1 |
| 124 | NDUFA5-628 | 0.603861 | 0.961038 | 165 | NDUFV1-1433 | 1 | 1 |
| 125 | UQCRC2-564 | 0.603861 | 0.636661 | 166 | NDUFV1-1687 | 1 | 1 |
| 126 | NDUFB10-637 | 0.604106 | 0.683456 | 167 | ND1-163-165 | 1 | 1 |
| 127 | NDUFS7-2083 | 0.608392 | 0.751295 | 168 | ND2-649 | 1 | 1 |
| 128 | COX4I1-312 | 0.613187 | 0.815099 | 169 | ND2-650 | 1 | 1 |
| 129 | NDUFS8-1164 | 0.614625 | 0.670986 | 170 | ND2-652 | 1 | 1 |
| 130 | COX7B-210 | 0.656184 | 0.941333 | 171 | COX4I1-2 | 1 | 1 |
| 131 | COX7B-208 | 0.656184 | 0.711797 | 172 | COX4I1-442 | 1 | 1 |
| 132 | NDUFA6-36 | 0.665217 | 0.811273 | 173 | COX4I1-443 | 1 | 1 |
| 133 | COX4I1-386 | 0.686753 | 0.729748 | 174 | COX6C2-139 | 1 | 1 |
| 134 | NDUFS8-101 | 0.688168 | 0.806344 | 175 | COX7A-4 | 1 | 1 |
| 135 | UQCRC2-3305 | 0.692795 | 0.754189 | 176 | COX7A-2188 | 1 | 1 |
| 136 | NDUFS7-2357 | 0.706331 | 0.964461 | 177 | ATP5F1B-5001 | 1 | 1 |
| 137 | NDUFV2-324 | 0.713982 | 0.78698 | 178 | ATP5F1C-1918 | 1 | 1 |
| 138 | COX7B-734 | 0.713982 | 0.925713 | 179 | ATP5F1C-2004 | 1 | 1 |
| 139 | ATP5F1C-1343 | 0.713982 | 0.925713 |  |  |  |  |
